# Proteomic Remodeling of the Cochlea During Chronic Suppurative Otitis Media Reveals Immune-Driven Injury Pathways

**DOI:** 10.64898/2026.02.27.707966

**Authors:** Ritwija Bhattacharya, Isha D Mehta, Viktoria Schiel, Jishnu Das, Vincent Yuan, Anping Xia, Peter L Santa Maria

## Abstract

Chronic suppurative otitis media (CSOM) is a persistent middle ear infection in which chronic inflammation drives irreversible sensorineural hearing loss (SNHL), yet the molecular mechanisms linking middle ear chronic infection to cochlear injury remain poorly defined. Using a murine model of CSOM induced by Pseudomonas aeruginosa (PA) persisters, a quantitative proteomic profiling of cochlear tissue was performed at seven days post-infection. Mass spectrometry revealed over 250 proteins that were significantly altered, indicating broad immune activation, oxidative stress, and disruption of ion homeostasis. Among the top candidates, HSPA5, MPO, and ATP2A2 were strongly associated with inflammatory stress and cellular injury pathways. These findings identify immunometabolic remodeling and macrophage-driven inflammation as central mechanisms linking chronic middle ear infection to cochlear injury.

## 1. Introduction

Chronic suppurative otitis media (CSOM) remains a major global health challenge and is the leading cause of hearing loss in children in low-resource settings [1,2]. It is characterized by chronic infection of the middle ear mucosa with persistent otorrhea, most commonly caused by Pseudomonas aeruginosa or Staphylococcus aureus [2,3]. Despite advances in antimicrobial therapy and surgical management, many patients with CSOM develop irreversible sensory hearing loss (SNHL), indicating that host-mediated inflammatory mechanisms play a critical role in disease progression beyond bacterial persistence alone [4].

Among CSOM pathogens, P. aeruginosa exhibits extraordinary persistence due to its ability to form biofilms and enter metabolically quiescent states that confer high tolerance to antibiotics and immune clearance[5]. Within biofilms, restricted nutrient availability, hypoxia, and altered ionic gradients induce adaptive resistance programs involving outer membrane remodeling, efflux pump activation, and metabolic rewiring [6,7]. These adaptations create a protected niche that perpetuates chronic infection and sustains prolonged immune activation within the middle ear cavity.

Our group established a murine model of CSOM that recapitulates the hallmark clinical and pathological features observed in humans, including persistent bacterial infection, tympanic membrane perforation, and progressive hearing loss [8]. Notably, this model revealed that cochlear dysfunction occurs in the absence of direct bacterial invasion, implicating immune-mediated bystander injury rather than pathogen dissemination. Instead, macrophages infiltrate the cochlea and mediate outer hair cell (OHC) loss, suggesting that immune-driven damage, rather than pathogen invasion, underlies auditory dysfunction [9]. However, the molecular triggers and signaling pathways linking chronic middle ear infection to cochlear immune activation remain unknown.

To address this gap, we employed an unbiased, nontargeted proteomics approach to delineate the cochlear proteomic landscape during CSOM. Using quantitative high-resolution mass spectrometry integrated with advanced bioinformatics, we systematically identified alterations in immune, metabolic, and stress-response pathways that distinguish infected from control cochleae. We therefore hypothesized that chronic middle ear infection induces a macrophage-driven immunometabolic stress program in the cochlea that primes immune-mediated injury prior to overt hair cell loss.

Collectively, our findings establish a mechanistic framework linking chronic inflammation in the middle ear to immune-mediated cochlear pathology. By defining the molecular basis of this cross-compartment immune crosstalk, our study identifies new therapeutic targets to mitigate hearing loss in CSOM and offers a foundation for a broader understanding of inflammation-driven sensorineural injury in the auditory system.

## 2. Materials and Methods

### 2.1. CSOM Model

#### 2.1.1. Animals and Ethics Approval

All animal experiments were conducted following approval from the Institutional Animal Care and Use Committee (IACUC) at Stanford University. Wild-type C57BL/6J mice (6–8 weeks old, JAX, Bar Harbor, ME, USA; Catalog #000664) were used in this study. Littermates were used for all experiments to ensure consistency. Animals were housed in a temperature and humidity-controlled facility at Stanford University’s animal care with ad libitum access to food and water. Anesthesia was used and maintained with ketamine (80–100 mg/kg) and xylazine (8–10 mg/kg) to minimize discomfort during all procedures [9].

#### 2.1.2. Preparation of PAO1 Persisters

A strain of PA with chromosomal integration of a luminescence reporter (PAO1.lux) was used for the experiment. For preparation, 20 µL of frozen PAO1 stock was added to 50 mL of Luria-Bertani (LB) broth and cultured overnight at 37 °C with shaking. The overnight culture was streaked onto an LB agar plate and incubated at 37 °C to isolate single colonies. A single colony was picked and inoculated into 10 mL of LB broth, followed by overnight incubation under aerobic conditions at 37 °C with shaking. The minimum inhibitory concentration (MIC) of ofloxacin against PAO1 was determined using a 96-well plate where the antibiotic was serially diluted in LB medium, and the bacteria were incubated at 37 °C overnight. Persister cells were generated by growing PAO1 in LB broth for 30 hours to reach the stationary phase. Ofloxacin was added at a concentration five times the MIC and incubated for another five hours. The bacterial culture was centrifuged, and the pellet was washed three times with phosphate-buffered saline (PBS) at 10,000 rpm for five minutes. The persister pellet was resuspended in PBS, and the concentration of persister cells was determined through serial dilution plating on LB agar. Colony-forming units (CFU) were counted after 48 hours, and a final concentration of 3.7 × 10⁶ CFU/mL was used in all experiments [10].

#### 2.1.3. Generating a CSOM Model

A validated murine model of CSOM was used in this study. Mice were anesthetized and positioned on a surgical stage under a microscope. Perforation of the tympanic membrane (TM) of the left ear was performed, followed by the inoculation of 5 µL of PAO1 persister cells (3.7 × 10⁶ CFU/mL) into the middle ear cavity. To maintain consistency, all inoculations were performed between 9:00 and 10:00 am.

Seven days post-inoculation, observation of the middle ear revealed signs of effusion, inflammation, and persistent TM perforation. Control mice underwent the same procedure but were inoculated with 5 µL of PBS into the middle ear cavity. Following inoculation, all mice were maintained in the same position with their left ear facing upward for 30 minutes. After 7 day, the cochleae were dissected and homogenized in protein lysis buffer for mass spectrometry studies [11].

#### 2.1.4. CSOM validation

To confirm CSOM, the middle ear of the mice was dissected and cultured on an LB agar plate for 48 hours. Mice were euthanized, and the middle ear tissues were harvested seven days following inoculation with persisters. Each sample was weighed and homogenized. The homogenate was then centrifuged at 1500 rpm for 5 minutes, and the resulting supernatant was collected. Serial dilutions of the supernatant were plated onto agar media, and colony-forming units (CFUs) were quantified after incubation for 48 hours. The CFU count was normalised to the weight of the middle ear tissue and expressed as CFU/ml per gram of tissue [10].

### 2.2. Method Details of Proteomics Analysis

#### 2.2.1. Sample Preparation for Mass Spectrometry

Six mouse cochlea lysates were treated with 1 μL Pierce™ Universal Nuclease (Thermo Scientific, San Jose CA). After vortexing and centrifugation, 25uL of each sample were transferred to new tubes and normalized to 50 μL with 10%SDS/100 mM TEAB for proteolytic digestion. The samples were reduced with 10 mM dithiothreitol (DTT) for 20 minutes at 55 degrees Celsius, cooled to room temperature, and then alkylated with 30 mM acrylamide for 30 minutes. They were then acidified to a pH ∼1 with 2.6 μL of 27% phosphoric acid and resuspended in 165 μL of S-trap loading buffer (90% methanol/10% 1M triethylammonium bicarbonate (TEAB)) and loaded onto S-trap microcolumns (Protifi, USA). After loading, the samples were washed sequentially with 150 μL increments of 90% methanol/10% 100mM TEAB, 90% methanol/10% 20 mM TEAB, and 90% methanol/10% 5mM TEAB solutions, respectively. Samples were digested at 47 degrees Celsius for two hours with 600 ng of mass spectrometry grade Trypsin/LysC mix (Promega, USA). The digested peptides were then eluted with two 35 μL increments of 0.2% formic acid in water and two more 40 μL increments of 80% acetonitrile with 0.2% formic acid in water. The four eluents were consolidated in 1.5 mL S-trap recovery tubes and dried via SpeedVac (Thermo Scientific, San Jose CA). Finally, the dried peptides were reconstituted in 2% acetonitrile with 0.1% formic acid in water for LC-MS analysis [12].

#### 2.2.2. LC-MS/MS Analysis

Proteolytically digested peptides were separated using an in-house pulled and packed reversed phase analytical column (∼25 cm in length, 100 microns of I.D.), with Dr. Maisch 1.9 micron C18 beads as the stationary phase. Separation was performed with an 80-minute reverse-phase gradient (2-45% B, followed by a high-B wash) on an Acquity M-Class UPLC system (Waters Corporation, Milford, MA) at a flow rate of 300 nL/min. Mobile Phase A was 0.2% formic acid in water, while Mobile Phase B was 0.2% formic acid in acetonitrile. Ions were formed by electrospray ionization and analyzed by an Orbitrap Exploris 480 mass spectrometer (Thermo Scientific, San Jose, CA). The mass spectrometer was operated in a data-dependent mode using HCD fragmentation for MS/MS spectra generation [12].

#### 2.2.3. Data Analysis

The RAW data were analyzed using Byonic v5.1.1 (Protein Metrics, Cupertino, CA) to identify peptides and infer proteins. FASTA files containing Uniprot *Mus musculus* and *pseudomonas aeruginosa* proteins and other likely contaminants and impurities were used to generate an *in silico* peptide library. Proteolysis with Trypsin/LysC was assumed to be fully specific. The precursor and fragment ion tolerances were set to 12 ppm. Cysteine modified with propionamide was set as a fixed modification in the search. Variable modifications included oxidation on methionine, dioxidation on tryptophan, glutamine and glutamic acid cyclization. Proteins were held to a false discovery rate of 1% using standard reverse-decoy technique [12].

### 2.3. Differential Protein Expression and Statistical Analysis

Protein abundance values were normalized and log₂-transformed prior to statistical analysis. Differential protein expression was assessed by comparing infected versus uninfected cochlear samples to identify proteins significantly altered in response to *P. aeruginosa* infection.

Statistical analyses were performed using appropriate computational tools (e.g., R). Proteins exhibiting statistically significant changes in abundance were identified using multiple testing correction, with significance defined based on Benjamini–Hochberg–adjusted p-values [13]. Results were visualized using volcano plots, displaying log₂ fold changes against adjusted significance values.

Principal component analysis (PCA) was conducted to evaluate global proteomic variation and assess clustering of samples by infection status. Hierarchical clustering and heatmap analyses were performed on significantly altered proteins to visualize expression patterns across biological replicates.

### 2.4. Pathway and Functional Enrichment Analysis

For functional enrichment analyses, identified proteins were mapped to their corresponding *Mus musculus* gene symbols prior to pathway analysis. This conversion enabled compatibility with gene-based annotation frameworks used for over-representation analysis (ORA) and gene set enrichment analysis (GSEA). All downstream enrichment analyses were therefore performed at the gene level, using mouse gene identifiers for pathway and Gene Ontology annotations. Enrichment analyses were performed using the clusterProfiler R package [14,15] to identify overrepresented pathways and functional categories.

Pathway annotations were derived from curated databases, including the Kyoto Encyclopedia of Genes and Genomes (KEGG) [16] and Reactome pathway [17] knowledgebase. Enriched pathways and Gene Ontology (GO) terms were identified based on adjusted p-values to account for multiple hypothesis testing, highlighting molecular processes associated with inflammation, immune activation, and cellular injury in CSOM-associated cochlear damage.

## 3. Results

### 3.1. CSOM Mouse Model

Persister cells (PCs) of P. aeruginosa were generated and inoculated into the mouse middle ear cavity to generate the CSOM model following the protocol of Xia et al. (2022) (Figure 1A). Xia et al. (2022) showed that there was significant outer hair cell (OHC) loss at 14 days after bacterial inoculation [9]. OHC loss was found predominantly in the basal turn (high-frequency region), with partial OHC loss in the middle turn (middle-frequency region) and no OHC loss in the apical turn (low-frequency region) of the cochlea. After examining OHC loss at three different timepoints: 3 days, 7 days, and 14 days, it was observed that OHC loss predominantly occurred in the basal turn of the cochlea at 14 days after bacterial inoculation. It was also established that macrophages are the major immune cells in the cochlea in CSOM, displaying increased numbers and distribution correlated with the observed cochlear injury. A significant elevation of macrophages in the CSOM cochlea was observed from day 3 of infection when compared to control cochleae (i.e., without infection), and this further escalated at 7 days and 14 days, showing a major increase in the number of macrophages (3 days vs control, P = 0.017; 7 days vs control, P < 0.001; 14 days vs control, P < 0.001)9. These findings indicate that macrophage accumulation precedes overt outer hair cell loss and identify the 7-day time point as a window to capture early immune and metabolic drivers of cochlear injury. Cochleae were therefore harvested at day 7 post-infection for proteomic analysis to identify early immune and stress-response pathways preceding structural damage (Figure 1B).

**Figure 1.**
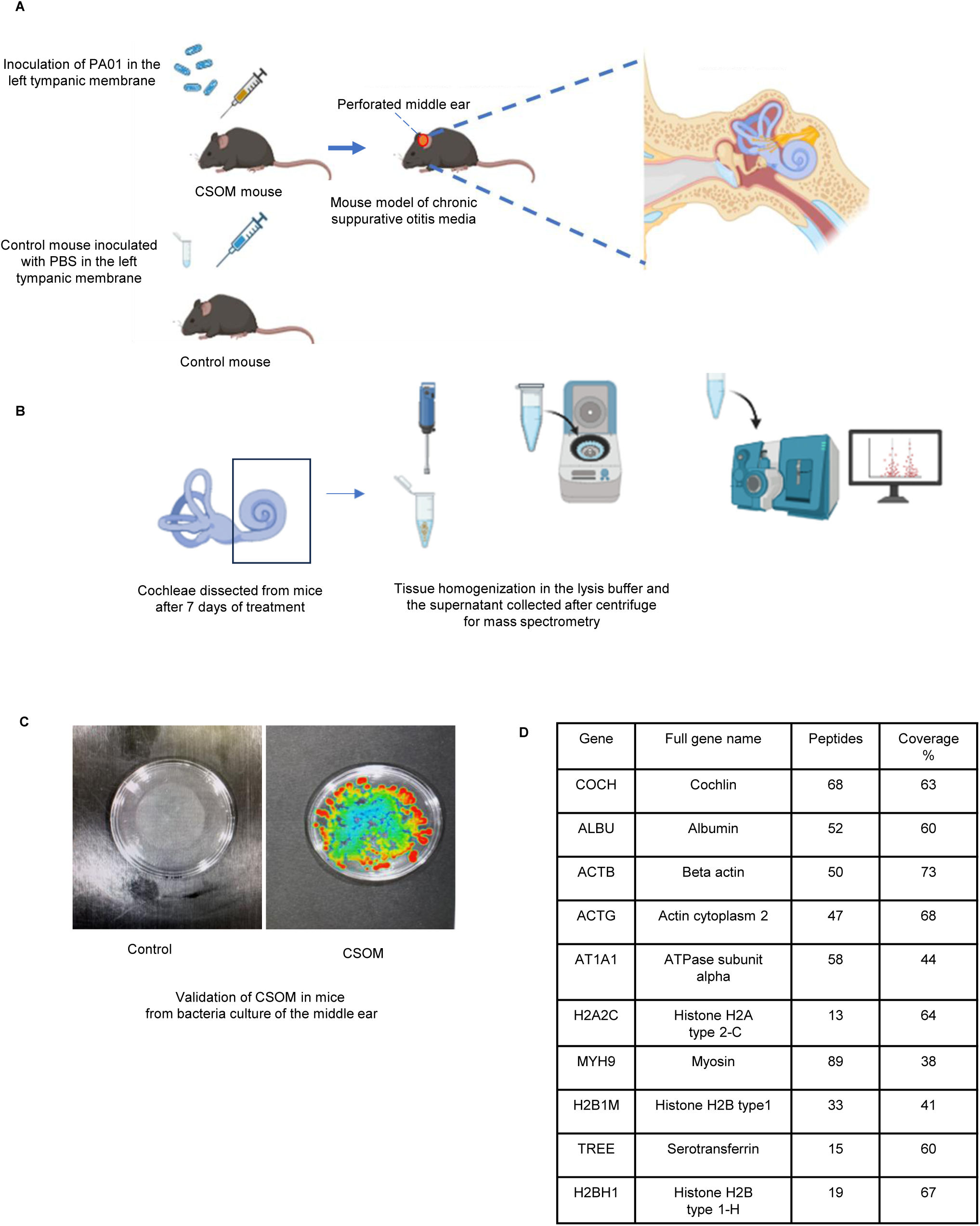
Experimental strategy for proteomic analysis of immune-mediated cochlear injury during CSOM. (A) Schematic of CSOM induction via inoculation of Pseudomonas aeruginosa PAO1 persister cells into the left middle ear following tympanic membrane perforation. Control mice received phosphate-buffered saline (PBS). The schematic illustrations in this figure were created with BioRender.com. (B) Experimental timeline illustrating cochlear harvest at day 7 post-infection to capture early immune and metabolic responses preceding overt hair cell loss. Cochleae were dissected, homogenized, and processed for quantitative mass spectrometry. The schematic illustrations in this figure were created with BioRender.com. (C) Representative bacterial culture images confirming persistent middle ear infection in CSOM mice, with no detectable growth in PBS-inoculated controls. (D) Summary of the top identified cochlear proteins detected by mass spectrometry, including Cochlin (COCH), Albumin (ALBU), and Actin isoforms, with corresponding peptide counts and sequence coverage.

To confirm the successful establishment of the CSOM model, bacteria were cultured from the middle ear tissues collected from both CSOM-induced and control mice. Middle ear homogenates were plated on Luria-Bertani (LB) agar containing cetrimide, a selective agent for Gram-negative bacteria. Bacterial growth was observed exclusively in the cultures derived from CSOM mice, whereas no colonies formed in control mice inoculated with phosphate-buffered saline (PBS). To further verify bacterial presence and viability, plates were imaged using an in vivo imaging system (IVIS). The luminescent signal, originating from genetically encoded bioluminescence in the bacterial strain, was clearly detected in the CSOM group, confirming successful colonization and persistence of the pathogen in the middle ear. These results collectively demonstrate the reliability of the CSOM mouse model used in this study (Figure 1, C).

PA modulates host responses through multiple, well-characterized mechanisms. In chronic infection settings, motility structures such as flagella and pili are often downregulated, reducing immune recognition. Alterations in flagellin expression can diminish Toll-like receptor 5 (TLR5) activation, and variations in lipopolysaccharide acylation can alter TLR4 signaling [18,19]. Additionally, Type II and Type III secretion systems deliver effectors that injure tissues and impair immune cell function; for example, ExoU can induce host-cell necrosis and disrupt inflammasome activity. Other factors, including pyocyanin and rhamnolipids, further compromise innate immune cells. These mechanisms collectively promote biofilm persistence, reduce bacterial clearance, and reflect the chronic, treatment-persistent course observed in clinical CSOM [2].

### 3.2. Global Proteomic Profiling of the CSOM Cochlea

Quantitative mass spectrometry of cochlear lysates from CSOM and control mice identified over 2,000 proteins with corresponding peptide counts and sequence coverage percentages. Figure 1D demonstrates the top 10 proteins identified in mass spectrometry.

Principal component analysis (PCA) demonstrated that PC1 accounted for 80% of total variance, robustly separating CSOM and control cochleae and indicating a dominant infection-driven proteomic shift (Figure 2A).

**Figure 2.**
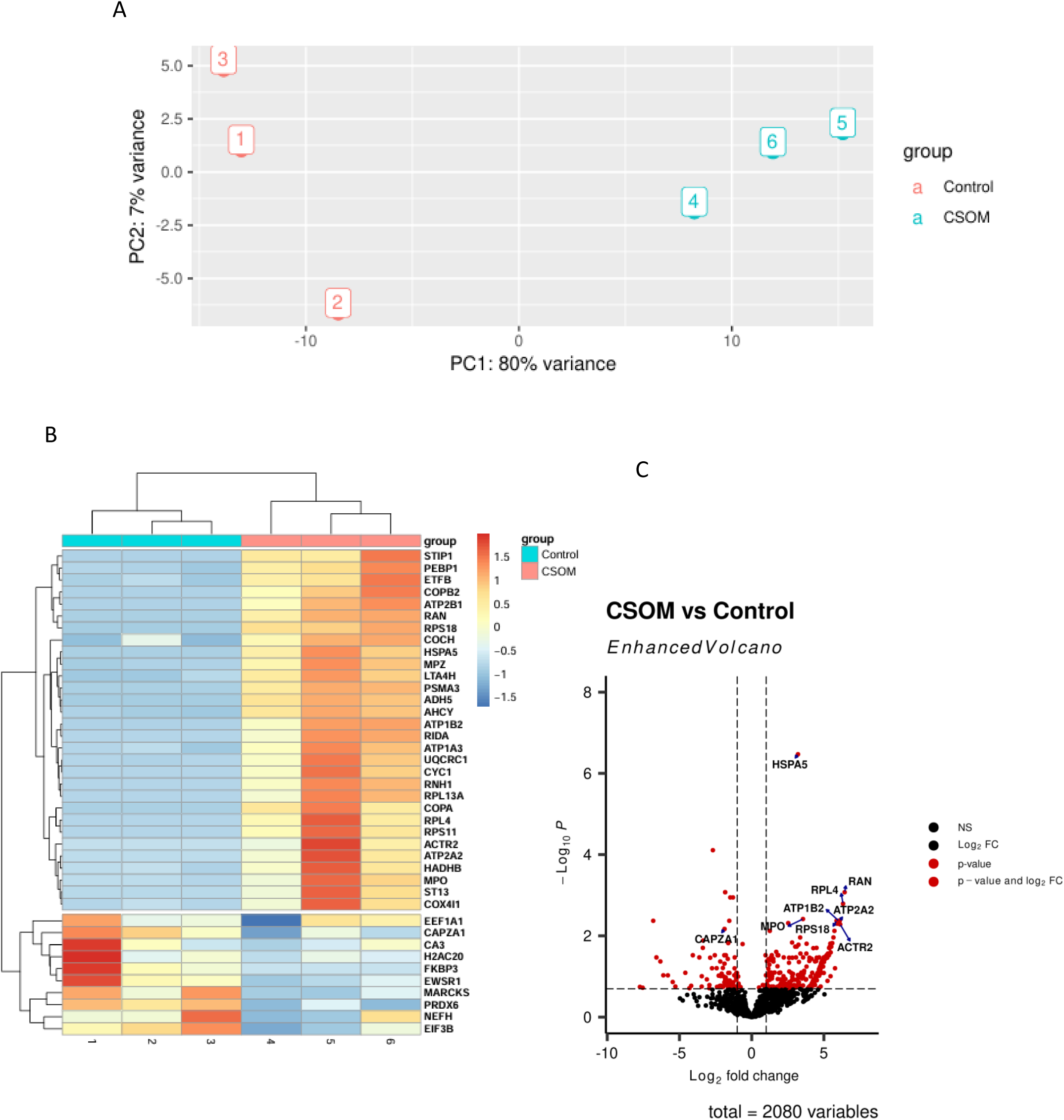
CSOM induces a distinct cochlear proteomic signature. (A) Principal component analysis (PCA) of cochlear proteomes from control and CSOM mice. PC1 accounts for 80% of total variance and robustly separates infected from control samples, indicating a dominant infection-driven proteomic shift. (B) Heatmap of the top 40 differentially expressed proteins across biological replicates. Relative protein abundance is shown, with red indicating increased and blue indicating decreased expression. Samples cluster by infection status. (C) Volcano plot depicting differential protein expression between CSOM and control cochleae. Proteins with significant upregulation or downregulation are shown based on log₂ fold change and adjusted p-value thresholds.

**Figure 3.**
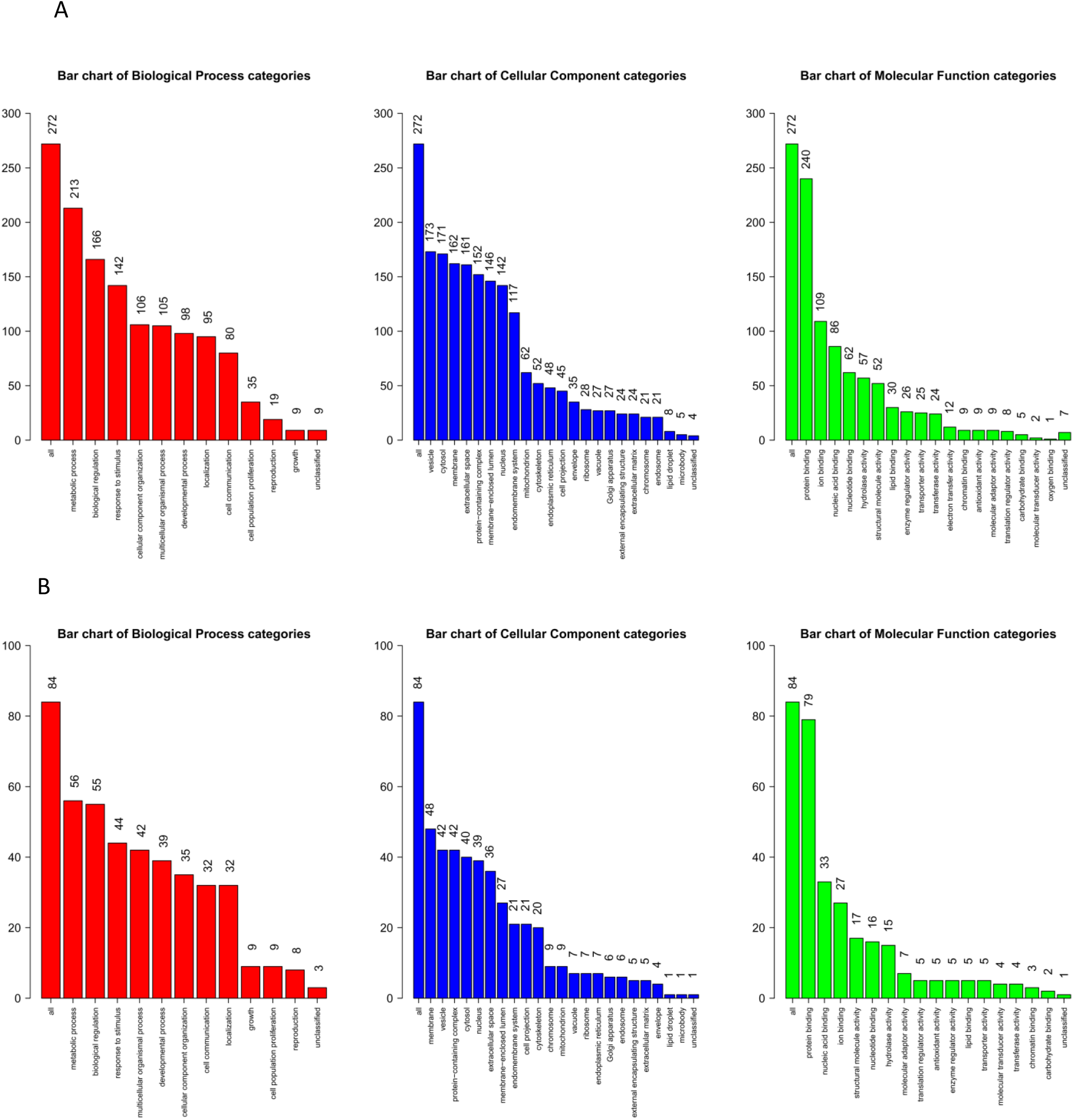
Gene Ontology enrichment reveals immune, metabolic, and stress-response pathways altered during CSOM. (A) Gene Ontology (GO) enrichment analysis of significantly upregulated proteins categorized by Biological Process, Cellular Component, and Molecular Function. (B) GO enrichment analysis of significantly downregulated proteins across the same functional domains. Bar plots indicate the number of proteins assigned to each GO category, highlighting coordinated remodeling of immune response, metabolic regulation, and cellular organization in the CSOM cochlea.

The heatmap shows differences in protein levels between control and CSOM cochlear samples. Samples from the control and CSOM groups cluster separately, indicating different protein expression patterns between the top 40 identified proteins in the two groups. Several proteins show higher expression in CSOM samples, while others show lower expression compared with controls. The color scale represents relative protein expression levels, with warmer colors indicating higher expression and cooler colors indicating lower expression (Figure2B).

To further illustrate these differences, a volcano plot was generated to visualise the distribution of significantly altered proteins (Figure2C). The plot displays the relationship between the log2 fold change and the -log10 adjusted p-value for all 2,080 quantified proteins, with proteins in the upper left and right quadrants representing significantly downregulated and upregulated proteins, respectively. Gene Ontology enrichment analyses of significantly altered proteins revealed coordinated changes in immune response, metabolic regulation, and cellular stress pathways (Figure3).

### 3.3. Candidate Genes Highlighted from Proteomic Analysis

Candidate proteins were prioritized based on statistical significance, magnitude of differential expression, and enrichment within immune- and metabolism-related pathways. Initially, proteins were ranked according to their fold change and corresponding p-values derived from the proteomic analysis. Genes demonstrating both high fold changes and strong statistical significance were prioritised. Subsequently, each candidate gene was cross-referenced with pathway enrichment analyses, including KEGG and Reactome databases (Figure 4). Only those genes that were significantly enriched within biologically relevant pathways highlighted in our analysis were retained. The final selection represents the candidate genes from the dataset, ensuring both statistical and functional significance. This analysis identified a core set of stress-response, inflammatory, and ion-handling proteins, including HSPA5, MPO, ATP2A2, and associated network components. These genes exhibited significant differential expression and were functionally enriched in pathways critical for protein folding and stress response (HSPA5), nucleocytoplasmic transport (RAN), ribosomal function and translational regulation (RPL4, RPS18), inflammatory response and oxidative metabolism (MPO, LTA4H), ion transport and calcium homeostasis (ATP1B2, ATP2A2), and cytoskeletal dynamics (ACTR2, CAPZA1) The significant role and functions of these genes and their associated pathways are validated using Uniprot database [20]. For instance, HSPA5 (also known as GRP78) plays a central role in endoplasmic reticulum (ER) stress response and protein quality control, processes that are often dysregulated during chronic inflammation. RAN, a small GTPase, is essential for nucleocytoplasmic transport, particularly during mitosis and cellular stress adaptation. MPO, a myeloid-derived enzyme, contributes to chronic inflammation through reactive oxygen species generation and has been implicated in immune-mediated tissue injury. Similarly, LTA4H mediates leukotriene B4 production, a potent inflammatory mediator^12^. ATP1B2 and ATP2A2 contribute to ionic homeostasis; ATP1B2 is a regulatory subunit of Na⁺/K⁺-ATPase involved in membrane potential maintenance, while the latter encodes SERCA2, a calcium pump regulating ER calcium levels. Both are crucial for epithelial integrity and cellular signaling. ACTR2 and CAPZA1, involved in actin polymerization and cytoskeletal organization, are essential for maintaining cellular morphology and motility under stress conditions. Ribosomal proteins RPL4 and RPS18 reflect translational reprogramming often observed during infection and immune modulation [20].

**Figure 4.**
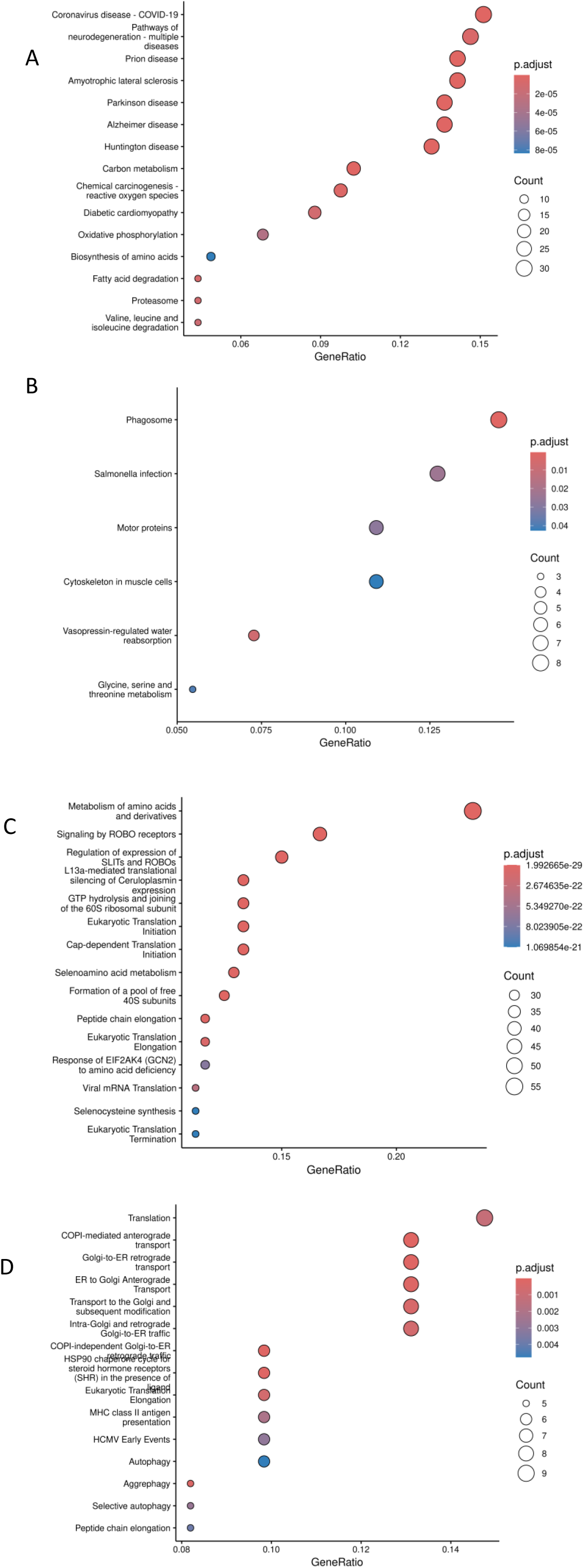
KEGG and Reactome pathway analyses reveal coordinated immune and metabolic pathway remodeling in CSOM. (A) KEGG pathways significantly enriched among upregulated proteins, reflecting shared immune, stress-response, and metabolic modules. (B) KEGG pathways significantly enriched among downregulated proteins, including pathways related to vesicular trafficking and cytoskeletal dynamics. (C) Reactome pathway enrichment analysis of upregulated proteins, highlighting amino acid metabolism, immune signaling, and translational initiation pathways. (D) Reactome pathway enrichment of downregulated proteins, including pathways involved in protein trafficking, autophagy, and translational regulation. Pathways are ranked by adjusted significance values.

**Figure 5.**
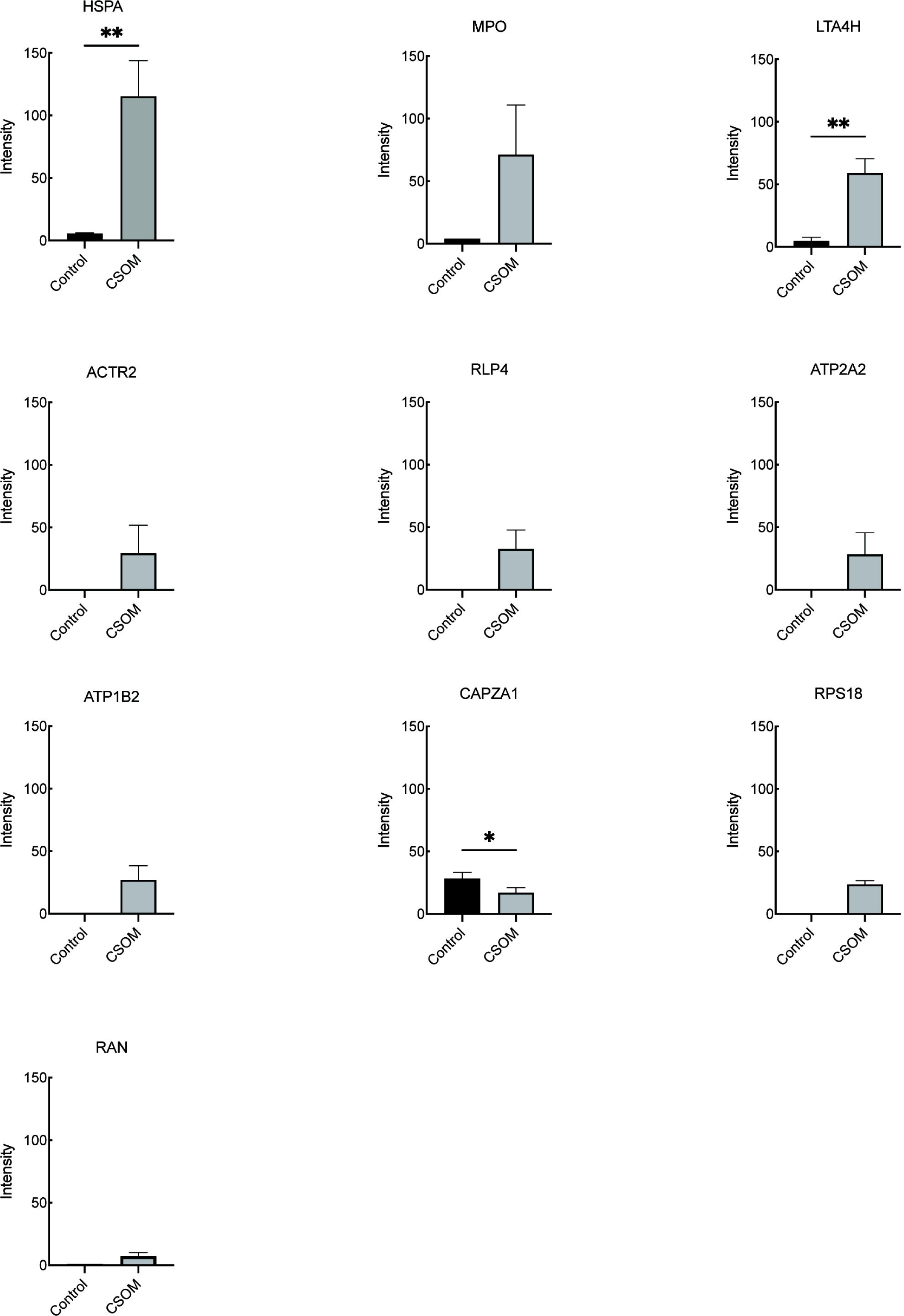
Identification of candidate immune and metabolic proteins associated with cochlear injury during CSOM. Relative abundance of selected candidate proteins identified through integrated differential expression and pathway enrichment analyses. Proteins involved in stress response, inflammation, and ion homeostasis, including HSPA5, MPO, ATP2A2, and LTA4H, are significantly upregulated in CSOM cochleae, whereas CAPZA1 is downregulated. Data represent mean ± SEM of mass spectrometry–derived intensity values (n=3 biological replicates per group). Statistical significance was determined by quantitative proteomic analysis (p < 0.05; p < 0.01).

Our findings are consistent with prior studies demonstrating macrophage infiltration into the cochlea during CSOM, where these cells drive outer hair cell loss [9]. Proteomic enrichment of stress-response chaperones such as *HSPA5* highlights activation of macrophages and engagement of inflammatory signaling pathways under chronic infection [21]. Elevated *MPO* suggests activation of myeloid cells that generate reactive oxygen species (ROS), driving collateral tissue injury [22–24]. Together, these changes define a distinct immunometabolic signature in which persistent infection induces macrophage activation, oxidative stress, and protein misfolding. This immune-driven stress environment likely primes cochlear tissues for delayed outer hair cell loss.

Among metabolic proteins, *ATP2A2* (SERCA2) and *ATP1B2* (Na⁺/K⁺-ATPase β2) were markedly dysregulated, indicating that chronic inflammation disrupts calcium and ion homeostasis in the cochlea [25]. Perturbation of ionic gradients and intracellular calcium handling can impair outer hair cell electromotility, driving progressive hearing loss even in the absence of direct microbial invasion and supporting a model of indirect inflammatory ototoxicity, where metabolic stress and cytokine signaling rather than infection itself mediate cochlear hair cell degeneration [26,27].

By integrating statistical significance, fold-change ranking, and pathway enrichment criteria, these ten genes provide a biologically meaningful snapshot of the proteomic alterations observed in CSOM. These targets not only underscore the mechanistic underpinnings of the condition under study but also offer promising candidates for further functional validation and potential therapeutic targeting.

## 4. Discussion

CSOM represents a paradigm in which chronic infection in one mucosal compartment drives immune-mediated injury in a spatially distinct sensory organ. *PA* enter the middle ear via a hole in the tympanic membrane and subsequently colonize and establish a biofilm community in the middle ear [9]. Currently, there is no effective medical cure to fully eradicate the infection. Biofilm-associated bacteria respond incompletely to antibiotic therapy, resulting in persistent immune activation despite microbiological suppression [28]. Current therapeutic options include topical antibiotics and antiseptics, as well as surgical procedures such as tympanoplasty or tympanomastoidectomy in refractory cases. While topical fluoroquinolones have the strongest evidence supporting their clinical use in CSOM, long-term treatment failure is common. These failures arise from the presence of persister cells, which are antibiotic-tolerant and capable of surviving treatment and acquiring antibiotic resistance [29]. Although povidone–iodine has been shown to clear both biofilm and planktonic cells in vitro assays, reports of ototoxicity limit its clinical utility [30]. The CSOM model exhibits prolonged persistence and shows poor response to fluoroquinolone antibiotics, as evident from previous studies [8]. A poor understanding of the mechanisms underlying PA-mediated sensorineural hearing loss (SHL) in CSOM hinders the development of a drug therapy to prevent hearing loss in hundreds of millions of children globally.

Our previous work established NLRP3 inflammasome activation in cochlear macrophages as a critical mediator of CSOM-associated hearing loss. It was established that the NLRP3 inflammasome has been implicated in various causes of hearing loss, including autoimmune disorders, tumours, and CSOM. It functions as an innate immune sensor primarily expressed in monocytes and macrophages. Upon activation, the inflammasome triggers a proteolytic cascade that culminates in the release of the proinflammatory cytokines IL-1β and IL-18 [10]. As observed by Xia et al. (2022), macrophages are the predominant immune cells present in the cochlea during CSOM. The progression of morphological changes further suggested a transition from monocytes into tissue macrophages. Consistent with these findings, cytokine activity in the CSOM cochlea reflected pronounced macrophage activation and inflammatory signaling. Macrophage elevation was shown to begin as early as day 3 and to increase substantially by days 7 and 14, correlating with the progression of cochlear injury, as OHC loss was predominantly and significantly found to occur at day 14 after bacterial inoculation into the mouse middle ear [9]. Therefore, our proteomics analysis at the 7-day timepoint provides important insight into the inflammatory and macrophage-associated pathways that likely precede and contribute to later OHC loss.

Several studies highlighted the association of the identified candidate genes in this study with the activation of NLRP3 pathways. In a recent study, Yang *et al.* (2025) reported that increased expression of HSPA5 in glioma promotes hypoxia tolerance and drives M2 macrophage polarisation, highlighting its role in shaping the inflammatory microenvironment. Similarly, in a different context, RPL4 has been associated with the enhanced release of damage-associated molecular patterns (DAMPs) [31,32]. DAMPs are endogenous molecules that amplify inflammation and, along with pathogen-associated molecular patterns (PAMPs) and recently defined homeostasis-altering molecular processes (HAMPs), these molecules are crucial in modulating innate immune responses. PAMPs and DAMPs are recognised intracellularly by cytoplasmic pattern recognition receptors (PRRs), the activation of which leads to inflammasome assembly and pyroptosis. Upon binding of a PAMP or DAMP, PRRs interact with apoptosis-associated speck-like protein (ASC), which oligomerises and uses its caspase activation and recruitment domain (CARD) to bind pro-caspase-1. The resulting PRR–ASC–pro-caspase-1 complex constitutes the inflammasome, where CARD–CARD interactions activate pro-caspase-1, leading to the maturation of pro-IL-1β and pro-IL-18 into their active, secreted forms (Brudette *et al.*, 2021) [33].

In addition, ATP1B2 upregulation in macrophages has been associated with oligodendritic damage through increased ferroptosis [21]. Ribosomal proteins, including RPS18, also exhibit potent bactericidal and agglutinating activity against both Gram-positive and Gram-negative bacteria [34]. Supporting this, a zebrafish model demonstrated that RPS18 recognises bacterial signature molecules such as peptidoglycan (PGN), lipopolysaccharide (LPS), and lipoteichoic acid (LTA), functioning as a pattern recognition receptor capable of binding and directly killing bacteria [35]. Moreover, a study identified LTA4H plays a pivotal role in macrophage aggregation and NF-κB activation [36]. Further, ACTR2 has been reported to promote activation of the NLRP3 inflammasome, pyroptosis, and inflammation in macrophages. Through paracrine effects, ACTR2-mediated IL-1β secretion also induces epithelial-mesenchymal transition (EMT) and fibrosis in tubular epithelial cells (TECs). Mechanistically, circACTR2 sponges miR-561, up-regulating NLRP3 expression and thereby enhancing IL-1β release, which in turn up-regulates fascin-1 and drives fibrosis [37].

Taken together, these findings collectively suggest that the molecular mediators identified in our proteomic analysis converge on shared inflammatory and macrophage-driven mechanisms that are central to NLRP3 inflammasome activation. In the context of CSOM, this alignment underscores a unified pathway in which chronic *PA* infection triggers macrophage activation and inflammasome-mediated cytokine release, ultimately contributing to cochlear inflammation and tissue injury. Collectively, these findings support a model in which chronic P. aeruginosa infection primes macrophage immunometabolic stress responses that converge on inflammasome activation, providing a mechanistic bridge between persistent infection and delayed cochlear injury.

## 5. Conclusion

This study delineates the immune and metabolic pathways that connect chronic *Pseudomonas aeruginosa* infection to cochlear injury in CSOM. Proteomic analysis seven days post-infection identified significant alterations in key proteins, including HSPA5, MPO, and ATP2A2, which map to inflammatory, immunometabolic, and cellular stress pathways. These molecular changes support a model in which macrophage-driven inflammation and immunometabolic stress converge to promote sustained cochlear damage. Together, our findings provide mechanistic evidence linking persistent infection to inflammation-mediated inner ear injury and highlight macrophage-centered pathways as potential therapeutic targets in CSOM-associated hearing loss.

## 6. Limitations of the study

Future integration of single-cell or spatial proteomic approaches will be essential to resolve cell-specific immune programs driving cochlear injury. Highly abundant proteins can mask those present at low abundance, and variable sensitivity compared with targeted approaches, such as Western blotting may have restricted the detection of low-level or transiently expressed proteins. Future studies using single-cell or spatial proteomic techniques could help overcome these limitations and better resolve the cellular sources and interactions driving cochlear inflammation.

## Declarations

### Ethics approval and consent to participate

All animal experiments were conducted following approval from the Institutional Animal Care and Use Committee (IACUC) at Stanford University.

### Consent for publication

All the authors agreed to the publication of this manuscript

### Availability of data and materials

The original datasets used during the current study are available from the corresponding author on reasonable request.

### Authors’ contributions

R.B, V.S and A.X did the experimental work. R.B, V.Y, I.M, J.D, and A.X performed or analyzed the immunological data computation. All authors contributed to the manuscript. PLSM, V.Y. led the manuscript writing and guided drafting. PLSM, AX planned the experimental designs.

## Acknowledgements

This work was supported by the Vincent Coates Foundation Mass Spectrometry Laboratory, Stanford University Mass Spectrometry (RRID:SCR_017801) utilizing the Thermo Exploris 480 nanoLC/MS system (RRID:SCR_022215). This work was supported in part by NIH P30 CA124435 utilizing the Stanford Cancer Institute Proteomics/Mass Spectrometry Shared Resource.

## Supplement information

**Supplemental information is available online**

---

**Figure S1. Generation and validation of *P. aeruginosa* PAO1 persister cells used for CSOM induction.**

(A) Schematic of PAO1 persister cell preparation, including bacterial culture, antibiotic treatment, and recovery.

(B) Confirmation of chromosomally encoded bioluminescence in PAO1 using in vivo imaging.

(C) Determination of the minimum inhibitory concentration (MIC) of ofloxacin for PAO1.

(D) Quantification of colony-forming units (CFU) across serial dilutions to establish inoculum concentration for middle ear infection.

