## Supplementary figures and images for "Proteomic Remodeling of the Cochlea During Chronic Suppurative Otitis Media Reveals Immune-Driven Injury Pathways"

### Supplementary figure 1

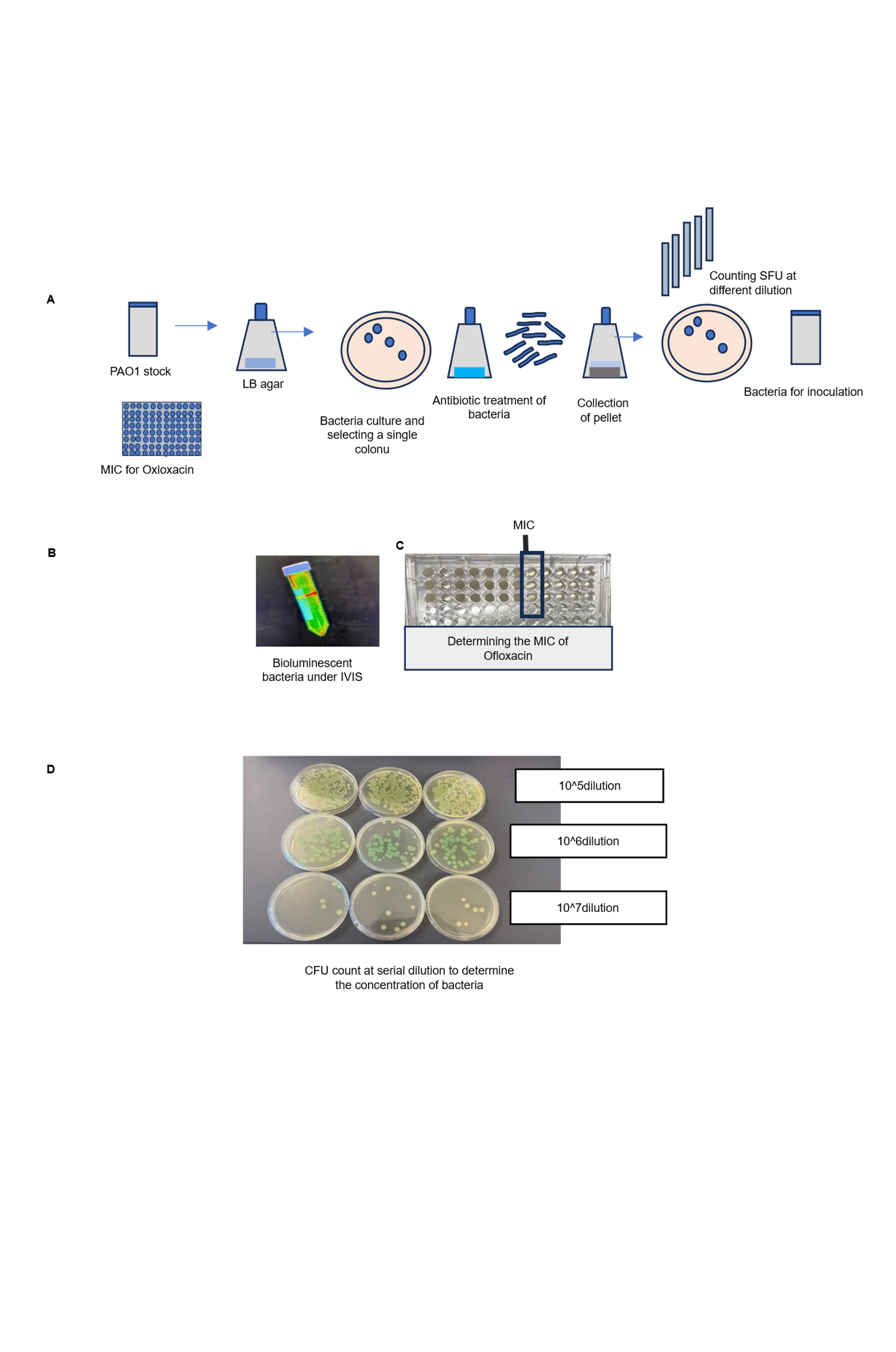
